# Disorganization impairs cognitive maps built from visual inputs

**DOI:** 10.64898/2025.12.22.695951

**Authors:** Xiaoyan Wu, Finn Rabe, Peizheng Wu, Victoria Edkins, Yves Pauli, Wolfgang Omlor, Akhil Ratan Misra, Stephanie Theves, Wolfram Hinzen, Iris E. Sommer, Philipp Homan

## Abstract

**Background and Hypothesis:** Cognitive disorganization is a core disturbance across the psychosis spectrum, characterized by difficulties integrating perceptual information into coherent representations. How such disturbances affect the construction of internal cognitive maps remains unclear. A key question is whether these maps emerge from visual input as readily as from language structure. We hypothesized that cognitive disorganization would impair cognitive maps derived from visual input while sparing performance when structure was provided linguistically.

**Study Design:** Participants with varying schizotypy learned food items associated with two attributes: who consumed the food (age group) and when it was consumed (time of day). These attributes were presented as images or language descriptions and organized in a two-dimensional (2D) conceptual space. Participants then completed similarity judgment and reward-learning tasks requiring inference of the underlying relational structure.

**Study Results:** Among individuals with higher cognitive disorganization, 2D distances were less predictive of similarity judgments, but only when relational structure had to be inferred from visual input. In the reward-learning task, computational modeling indicated that decisions were guided by spatial representations (Gaussian process). Higher cognitive disorganization was associated with weakened spatial generalization when structure had to be inferred from visual input, but not when it was provided through language.

**Conclusions:** Cognitive disorganization disrupts the ability to build and use cognitive maps from visual input, while performance appeared preserved when structure was conveyed through language. These findings support the shallow cognitive map hypothesis and suggest that cognitive disorganization selectively disrupts the transformation of perceptual input into stable relational representations.

## Introduction

When exploring a new city, we integrate perceptual fragments such as streets and landmarks into an internal representation that supports navigation in the city. A similar representation might also be formed without direct experience; for instance, one may gain an impression of a city’s layout solely from a friend’s vivid written description. This may be because linguistic descriptions activate pre-existing knowledge, enabling the formation of an internal structure even without direct perceptual experience. Understanding how such fragmented experiences are organized into a coherent internal structure is central to theories of memory, reasoning and prediction. A widely held view is that the hippocampal-entorhinal system constructs cognitive maps that encode relationships among concepts or events, enabling flexible generalization across physical and abstract spaces^1–5^. Recent work has shown that this mapping system is also relevant to general cognitive ability^6,7^. Cognitive maps provide a framework that supports the inference of latent relational structure^8^, the prediction of expected outcomes^9^ and value-guided choice^10,11^.

The brain mechanisms involved in the development and maintenance of cognitive maps appear to be particularly vulnerable in schizophrenia spectrum disorders. A core psychosis spectrum trait, cognitive disorganization, is characterized by difficulties in organizing information and maintaining coherent cognition^12^. In individuals with schizophrenia, language output disturbances are widely considered to reflect disorganized thinking and abnormalities in conceptual and semantic organization (for review, see^13^). The recently proposed shallow cognitive map hypothesis suggests that cognitive disturbances in schizophrenia may reflect disruptions in the hippocampal-entorhinal system that supports relational structure^14,15^. According to this hypothesis, hippocampal-entorhinal attractor dynamics become less stable, yielding shallower attractor basins and undermining coherent relational structure. Consistent with this hypothesis, individuals with schizophrenia show reduced reliance on semantic proximity when generating words within a category, and greater reliance on semantic structure has been linked to stronger replay-associated ripple power measured during a post-task rest session^16^. Individuals with schizophrenia also exhibit impaired accuracy in physical-space navigation and diminished grid-like signals in the entorhinal-hippocampal system^17^.

However, it remains unclear whether the cognitive disorganization characteristic of the psychosis spectrum reflects a generalized failure in relational reasoning or a specific vulnerability in transforming fragmented perceptual inputs into coherent internal structures. A recent study^17^ showed that schizophrenia is associated with impairments in physical spatial navigation, but the impact on the navigation of the relational structure derived from abstract conceptual knowledge remains untested. Studies of abstract knowledge mapping have typically used visual stimuli such as images or scenes as the inputs to dimensional spaces^5,10,11,18,19^, which provide no semantically explicit information. In these paradigms, the mapping between objects and their locations in the space is often arbitrary, for example, pairing the length of a duck’s neck and legs with a Christmas icon^18^.

Other studies used audiovisual stimuli with newly learned symbolic labels, and similar grid-like neural coding has also been observed for the resulting two-dimensional (2D) feature spaces^20, 21^. However, the associations in these studies lack long-term semantic representations and thus do not address whether similar coding principles apply to real-world word meanings. Consequently, the role of explicit semantics, such as words or texts, in facilitating internal relational structure, relative to purely visual inputs, remains less explicitly investigated. No study has directly compared these two forms of input and examined whether they differentially affect individual differences associated with the psychosis spectrum.

Schizotypal traits are widely considered a dimensional expression of psycho-sis-related vulnerability in the general population and are commonly used to investigate variability in nonclinical samples. In the present study, schizotypal traits were measured using the Oxford-Liverpool Inventory of Feelings and Experiences (O-LIFE)^22^. Here we examine how individual differences in cognitive disorganization influence the construction and use of cognitive maps derived from fragmented inputs presented as image-based or language-based stimuli. Participants learned two graded dimensions by fragmented exposure, associated them with food items, and subsequently used the inferred 2D conceptual map to support similarity judgments and re-ward-based decisions. We found that when the inputs were image-based, higher cognitive disorganization was associated with reduced spatial sensitivity in the similarity judgments, and reduced spatial generalization in reward-based decisions. No such association was observed when the inputs were language-based. These findings suggest that a central vulnerability in the psychosis spectrum lies in transforming visual inputs into a stable internal structure, whereas the ability to operate on already organized, language-based structure appeared relatively preserved.

## Methods

This study is a pre-registered study (https://aspredicted.org/z8tn-kjvw.pdf), and all analyses followed the prespecified plan unless otherwise noted. Fisher’s z-tests for between-group comparisons and Bayes factor analyses for non-significant results were conducted as additional analyses to quantify evidence and were not included in the original pre-registration.

### Participants

Our target sample size was 80 participants. Given that the study involved demanding training tasks, we recruited more volunteers to compensate for potential dropouts or incomplete sessions. The final sample had a total of one hundred and eleven participants, with 51 in the image group and 60 in the language group. Participants were right-handed adults (≥18 years) recruited via social media, with normal or corrected-to-normal vision, C1-level or higher English proficiency, and no prior education or training in psychology. The final sample ranged in age from 18 to 64 years. Written informed consent was obtained from all participants prior to the experiment. This study adhered to the principles of the Declaration of Helsinki and was approved by the Cantonal Ethics Committee Zurich (Kantonale Ethikkommission Zürich; Protocol No. 2024-01314).

### Experiment procedures

Participants were assigned to one of two between-group input conditions. The grading of each of the two variables (i.e., age group and time of day) were presented as images with symbolic icons in the image-cued condition and as English text in the language-cued condition. The whole experiment included several parts and lasted 2-3 hours (mean = 2 h 10 min; Fig. 1a).

**Figure 1:**
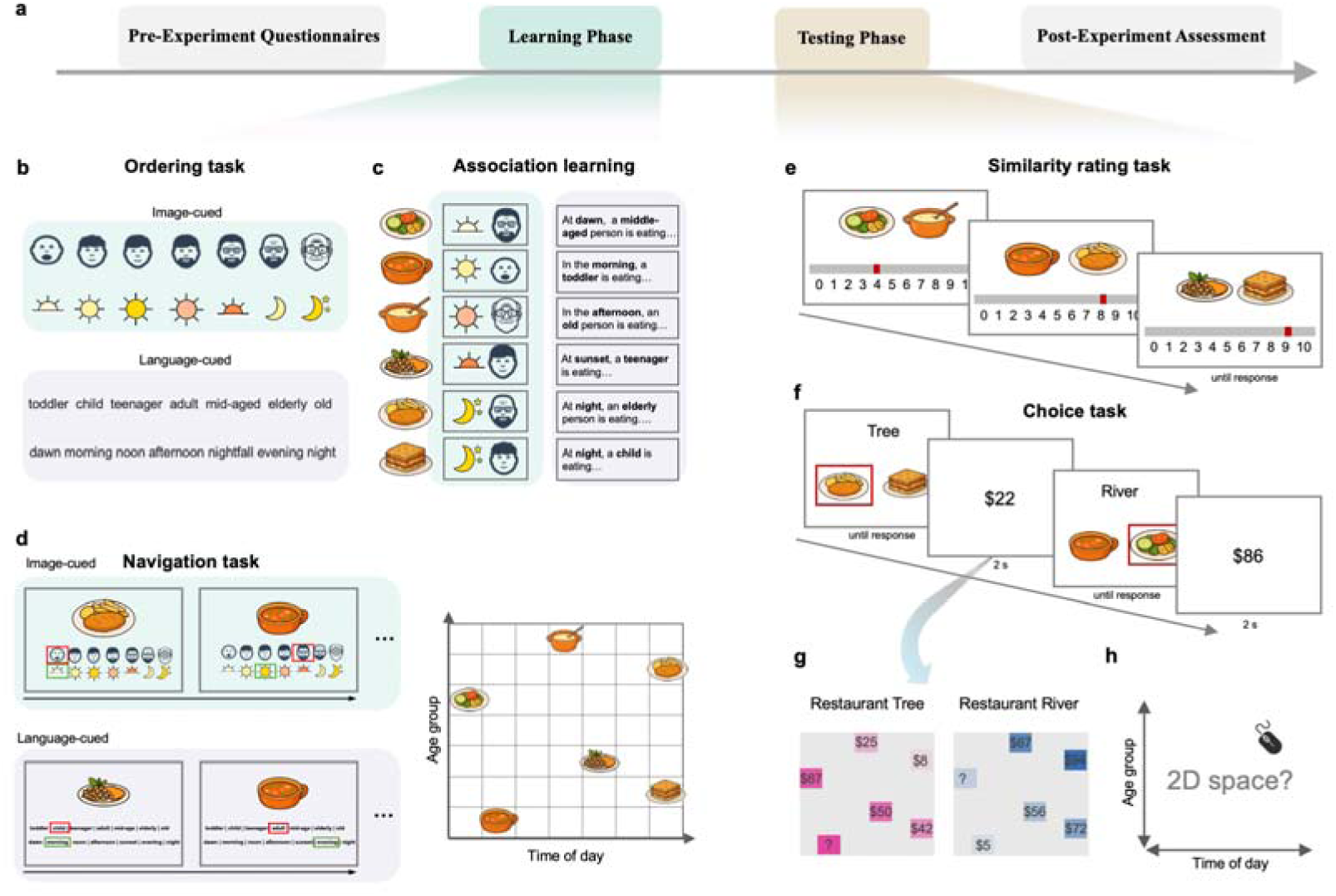
Overview of the experimental procedure and tasks. **a.** Schematic timeline of the experiment, including the pre-experiment questionnaires, the learning phase, the testing phase, and the post-experiment assessment. **b.** Ordering task: participants learned the grading of two variables, the age group and time of day, presented either with images (upper panel: faces and sun-moon icons; image-cued condition) or with English words (lower panel: language-cued condition). **c.** Association learning: participants memorized six food items with a specific combination of age group and time-of-day cues. The stimuli of two grading variables were either presented as images (left panel: image-cued condition) or English short sentences (right panel: language-cued condition). **d.** Navigation task: participants were shown a food item and asked to select the correct “age group” and “time of day” associated with it. Each trial required participants to make both selections correctly. **e.** Similarity rating task: participants rated the similarity between pairs of food items on a 0-10 scale based on their learned age-time associations. **f.** Choice task: participants chose between two food items on each trial. After each decision, the value of the selected item was displayed as feedback. Food items were drawn from either the Tree or River “restaurant,” with their values determined by their locations in an unshown 2D space defined by age group and time of day. **g.** Value structures used in the choice task. Each participant completed the task in two value contexts (Restaurant Tree in pink, Restaurant River in blue), with context-value structure assignment randomized across participants. Values were generated from item coordinates using either 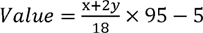 (pink structure) or 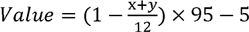 (blue structure). Items marked with question marks were never shown in the choice task. **h.** Placement task: participants were shown a 2D map and asked to drag each food item to its estimated location within the map.

### Pre-experiment questionnaires

Participants provided demographic information and completed standardized personality measures, including the short form of the *0*−*LIFE* for schizotypy^22^, the Big Five Inventory^39^, and the Positive and Negative Affect Schedule (*PANAS*)^40^. To account for potential appetite-related confounds arising from the food images, participants also reported their hunger level and the time since their last meal. Results are reported in Supplementary Table S1.

### Learning phase

In the learning phase, participants memorized each food item together with its corresponding “features,” namely who ate it (age group) and when it was eaten (time of day). Participants were required to learn the grading of each of the two variables, presented either as image-based or language-based stimuli, through sequential fragmented exposure. They then learned the associations between the graded variables and the food items. No terms referring to “maps” or “two-dimensional spaces” were ever presented. Previous research has shown that such two-dimensional association learning can lead to the formation of an internal two-dimensional representational space, i.e., a cognitive map^3,4^.

### Ordering task

Participants learned the grading of two variables, age group and time of day, through sequential fragmented exposure. Each variable consisted of seven graded levels (age group: toddler, child, adolescent, adult, middle-aged, elderly, old; time of day: dawn, morning, noon, afternoon, nightfall, evening, night). In each trial, participants judged the relative order of two randomly selected levels from the same variable (e.g., whether “toddler” is younger or older than “adolescent”). Training continued until participants reached at least 85% accuracy over the most recent 20 trials, ensuring that they understood the ordinal structure of each variable. The learning order of the two variables was randomized across participants. After reaching the accuracy criterion, participants were asked to arrange the seven levels of each variable from smallest to largest. Note that the inputs for the two variables were presented using different types of stimuli in the image-cued and language-cued groups. In the im-age-cued group, the categories were represented by cartoon-style faces (age) and symbolic icons (time of day, e.g., a sun or moon), whereas in the language-cued group they were presented as English words (Fig. 1b).

### Association learning

Participants then learned the associations between the graded variables and the food items. The task involved six food items paired with specific combinations of age group and time of day. In each trial, a target age-time combination was shown at the top of the screen along with six candidate food items below. Participants selected the correct food item using the mouse and received immediate feedback on accuracy. Training continued until participants reached at least 85% accuracy across the most recent 20 trials. Again, the inputs for the age-time combinations were presented using different types of stimuli in the image-cued and language-cued groups (Fig. 1c).

### Navigation task

Participants were presented with a food item and asked to identify the corresponding time of day and age group. In each trial, a target food item was shown at the top of the screen. Participants selected the correct age group and time of day from all graded levels. Feedback was provided immediately after each selection, and a trial was recorded as correct only if both the time and age choices were correct on the first attempt. Training continued until participants reached 12 consecutive correct trials. Again, all possible options covering the full grading range were presented using different stimulus formats in the image-cued and language-cued conditions (Fig. 1d). Performance accuracy in the above tasks did not differ significantly between the two conditions (see Supplementary Fig. S1).

### Testing phase

Following the learning phase, participants proceeded to the testing phase.

### Similarity rating task

Participants judged the similarity of food-item pairs based solely on the learned graded variables (Fig. 1e). In each trial, two food images appeared side by side, and participants rated their similarity from 0 (not similar) to 10 (highly similar) using a mouse-controlled slider with a random starting position. The task comprised 60 trials. Participants were explicitly instructed to judge the similarity between pairs of food items based solely on who ate them and when they were eaten, and not on any other attributes such as taste, color, or prior pairings.

### Choice task

Participants then completed a choice task in which unknown rewards depended on each item’s location within an unshown 2D map defined by the learned graded variables (Fig. 1f). Participants were given a cover story stating that the price of each food item depended on both the eater’s age group and the time of day at which it was eaten, and that the way in which these two factors affect price differs between the two restaurants, Tree and River. Unbeknownst to participants, the two restaurants followed different location-value mappings (Fig. 1g), and the specific mapping rules were not disclosed. The exact pricing computation is provided in the caption of (Fig. 1). Note that each price included a ± 2 noise term. The assignment between restaurant names (Tree, River) and the corresponding value maps was randomized across participants.

The task consisted of blocks associated with each of the two restaurants. At the beginning of each block, participants were informed which restaurant they were in, and a Tree or River symbol remained at the top of the screen throughout that block. In each trial, two food items appeared side by side, and participants selected one using the left or right arrow key on a keyboard. Only the value of the chosen item was revealed after making a choice. Participants were instructed to maximize their rewards. The task consisted of four blocks (two for each restaurant) with 30 trials per block, resulting in 120 trials in total.

At the end of the choice task, participants were asked to estimate the values of all food items for each restaurant based on what they had experienced during the task. Unbeknownst to participants, the estimation phase included one additional food item per restaurant (i.e., the question mark in Fig. 1g) that had never been presented during the choice task. This manipulation allowed us to assess the model’s ability to generalize value estimates to untrained items.

### Placement task

After the choice task, participants were asked to place the food items onto a 2D map defined by age group and time of day. The grid was structured by age (youngest to oldest; vertical axis) and time of day (earliest to latest; horizontal axis), as shown in Fig. 1h. All food items initially appeared outside the grid and participants used the mouse to drag and drop each item onto the location corresponding to its previously learned age-time association.

### Post-experiment assessment

After the experiment, participants completed a brief questionnaire assessing whether they experienced any uncertainties during the task and whether such uncertainties influenced their choices. They also performed two additional cognitive tasks: an N-back task^41^ to index working memory capacity and a mental rotation task^42^ to assess spatial transformation ability, each lasting approximately 7-8 minutes. In the final session, participants completed an audio recording task; these data were not included in the present analyses.

### Behavioral analysis

As specified in the pre-registration, similarity ratings were analyzed using a linear mixed-effects model (LMM) with similarity as the dependent variable. Fixed effects were group (image vs. language), 2D distance, and their interaction, with random intercepts for participants. The model was fitted using maximum likelihood estimation (default) in MATLAB (fitlme). All participants were included in this analysis.

### Computational modeling

As specified in the pre-registration, we compared four computational models with different hypotheses about decision-making during the choice task to examine how participants learned and generalized rewards across contexts. The data from all participants were included in this analysis.

### Model 3: Spatial gaussian process regression (GPR) model

Participants’ behavioral choices were best described by Model 3. Details of alternative models (Models 1, 2, and 4), model fitting, as well as comparison and validation, are provided in Supplementary methods.

This model assumes that participants generalize experienced rewards across the 2D spatial layout of the map using Gaussian process regression (GPR). Each map context is modeled independently, maintaining its own history of chosen locations *X* = {*x*1,…, *xn*} and corresponding rewards *Y* = (*r*1,…, *rn*) ⊤. The covariance between any two past choices is computed via a radial basis function (RBF) kernel:

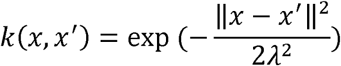

where *λ* is the lengthscale parameter governing the extent to which input similarity decreases with distance, reflecting the weight assigned to spatial information in the generalization process. Larger *λ* values produce broader generalization across space.

For each candidate option *x*, the predicted mean reward is

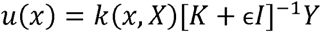

where *K* = *k* (*X, X*) denotes the kernel matrix whose (*i,J*)-th entry is given by *K_i j_* = *k* (*x_i_*, *x_j_*). A small jitter term □*I*, where *I* denotes the identity matrix, is added to the kernel matrix to ensure numerical stability when computing its inverse. Choices follow a softmax over the predicted means:

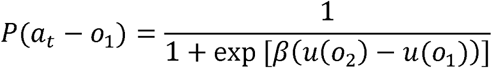

Free parameters: *λ* (spatial lengthscale) and *β* (inverse temperature). o_1_ and o_2_ denote Option 1 (shown on the left) and Option 2 (shown on the right), respectively.

## Results

The experimental procedure included a learning phase and a testing phase (Fig. 1a). During the learning phase, participants learned six food items and their associated attributes along two abstract dimensions: the age group of the person who eats the food (seven ordered levels ranging from toddler to old) and the time of day of eating (seven ordered levels ranging from dawn to night). These two dimensions formed a 2D conceptual map. During training, participants were randomly assigned to either an image-cued (*N* = 51, Fig. 1b-d, green-shaded panels) or a language-cued (*N* = 60, Fig. 1b-d, gray-shaded panels) condition, in which stimuli representing the two dimensions were presented in the form of images or text, respectively.

Participants first learned the rank relationships within each dimension by judging the relative order of two levels (e.g., whether “toddler” is younger or older than “adolescent”) until performance achieved an accuracy threshold (85%). They then learned the six associations of food attributes through association learning (Fig. 1c), in which the participants selected the correct food item given a specific combination of attributes, and navigation tasks (Fig. 1d) that required identifying the correct attributes for each food item. Training continued until the performance reached the same threshold.

In the subsequent testing phase, we assessed participants’ ability to infer unobserved relationships within the 2D conceptual map using a similarity rating task (Fig. 1e), as well as their ability to use map-based information to support reward generalization using a choice task (Fig. 1f). All participants completed identical tasks and were presented with the same stimuli during the testing phase. All participants completed the experiment in a single session conducted on the same day.

### Higher cognitive disorganization reduces spatial sensitivity specifically in visually derived cognitive maps

We first examined whether distances on the unshown 2D map (Fig. 1d, right panel) could predict participants’ judgments of similarity between food items (Fig. 1e), and how cognitive disorganization (assessed using the OLIFE) influences this process. Participants were instructed to judge the similarity between pairs of food items based solely on who ate them and when they were eaten.

Beyond the preregistered analyses, a linear mixed-effects model revealed that dissimilarity scores (reverse-scored similarity) increased with 2D distance (*β* = 0.87, 95% CI [0.78, 0.96], *p* < 0.001), and that the interaction between distance and group was significant (*β* = −0.31, 95% CI [−0.44, −0.18], *p* < 0.001; Fig. 2b). We further observed significant interactions involving cognitive disorganization, including a three-way interaction between distance, cognitive disorganization, and group (see Supplementary Table S2). These results suggest that participants were able to infer unobserved relationships within the 2D conceptual map, despite never seeing the map directly, but this effect differed between the image and language groups. We also found the 2D-distance model outperformed alternative 1D models (see Supplementary Analysis for details).

**Figure 2:**
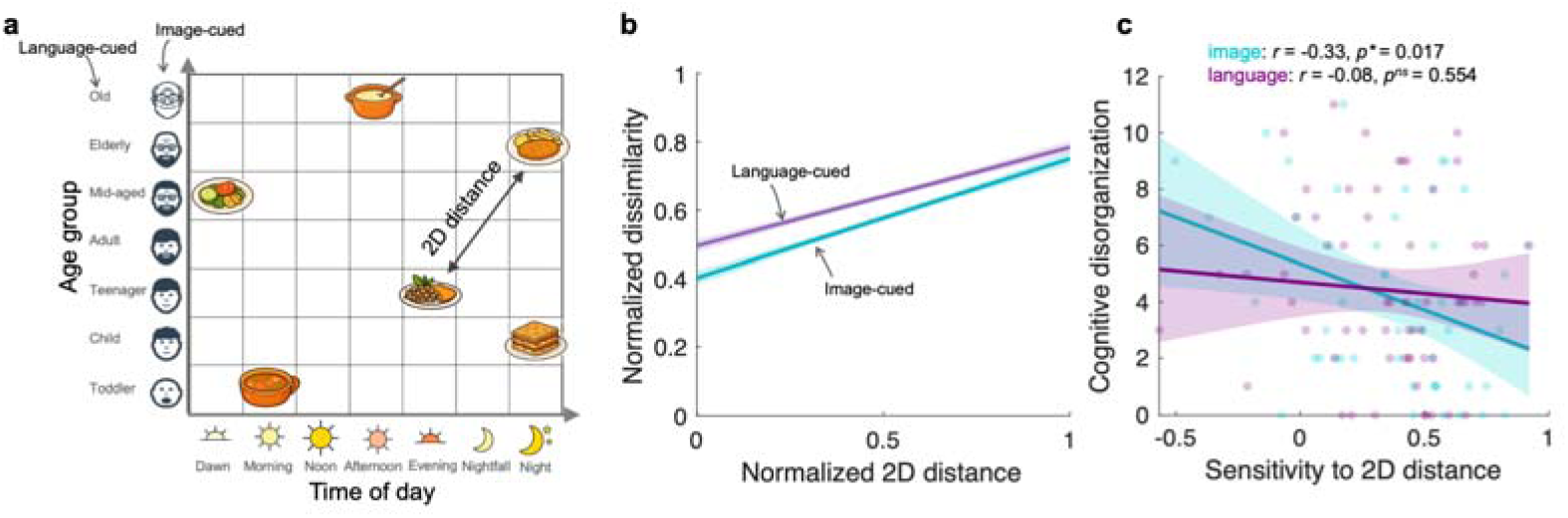
Results of the similarity rating task. **a.** Example of the 2D age-time abstract space used to define pairwise distances between food items. The solid arrow illustrates the 2D Euclidean distance. **b.** Results of the similarity rating task. The dissimilarity scores (computed as 10 - similarity), min-max normalized to the [0, 1] range, increased with spatial distance in both the image-based and language-based conditions. **c.** Correlation between cognitive disorganization scores (O-LIFE subscale) and sensitivity to 2D distance. Sensitivity to 2D distance was quantified as the Fisher z-transformed correlation between pairwise 2D distances and the corresponding dissimilarity scores, with higher values reflecting stronger spatial sensitivity. For panels b and c, lines show separate linear fits for the image (cyan) and language (purple) conditions, with shaded areas representing 95% confidence intervals. Significance levels: *p < 0.05; ns = not significant.

We next examined whether the ability to infer relationships within the 2D conceptual map was associated with cognitive disorganization. Here the spatial sensitivity was quantified as the Fisher’s z-transformed correlation between pairwise 2D distances and dissimilarity scores. We found that spatial sensitivity was negatively correlated with cognitive disorganization in the image-cued condition (*r* = −0.33, 95%*CI* [−0.56, −0.06], *p* = 0.017), but not in the language-cued condition (*r* = −0.08, 95%*CI* [−0.33, 0.18], *p* = 0.554, *BF*_01_ = 6.45; *Fis*□*er′sZ* = −1.34, *p* = 0.090; Fig. 2c).

A similar dissociation between conditions was also observed for the association between spatial sensitivity and the total O-LIFE score: a significant correlation was observed in the image-cued condition but not in the language-cued condition (see Supplementary Fig. S3a). In contrast, other personality measures, such as the Big Five, showed no associations with spatial sensitivity (Supplementary Fig. S3 and Fig. S6), indicating that the link between spatial sensitivity and schizotypy-related cognitive disorganization is selective rather than reflecting general personality variation.

Furthermore, in the image-cued condition, sensitivity to 2D distance was positively correlated with mental rotation performance (*r* = 0.34, 95%*CI* [0.07, 0.56], *p* = 0.015), linking spatial sensitivity to spatial transformation ability. No such relationship was observed in the language-cued condition (*r* = 0.02, 95%*CI* [−0.24, 0.27], *p* = 0.908, *BF*_01_ = 7.69; *Fis*□*er′sZ* = 1.71, *p* = 0.044; see also Supplementary Fig. S6).

In brief, participants’ similarity judgments reflected distances within an unobserved 2D conceptual map. In the image-cued condition, higher levels of cognitive disorganization were associated with lower spatial sensitivity, and higher spatial sensitivity was associated with better spatial transformation ability. These associations were not observed in the language-cued condition.

### Spatial memory predicts reward learning performance specifically in visually derived cognitive maps

If participants form an internal spatial representation from sequentially fragmented inputs, better memory for this space should support more effective use of spatial information for problem solving. To this end, participants completed a choice task in which they selected one of two food items and received a reward for the chosen option (Fig. 1f). They were instructed to choose the option with the higher reward. Unknown to participants, the rewards associated with the food items were determined by linear functions of their locations in the 2D space (Fig. 1g).

We found that participants in both the image-cued and language-cued conditions improved their decisions over time (Fig. 3a). At the end of the experiment, participants were asked to locate all food items within a 2D space in a placement task (Fig. 1h). Accuracy was quantified as the distance between participants’ placed locations and the true coordinates, defined as the 2D placement error (Fig. 3b&3c). Participants were generally able to place the food items close to their correct locations, and no difference was observed between the image-cued and language-cued conditions (*t* (109) = 0.79, 95%*CI* [−0.68, 1.57], *p* = 0.432, *BF*_01_ = 7.68; Fig. 3d). These results indicate that, despite never seeing the 2D map directly and reporting no explicit awareness of a map-based strategy, participants were nonetheless able to reconstruct the hidden spatial structure with considerable accuracy.

**Figure 3:**
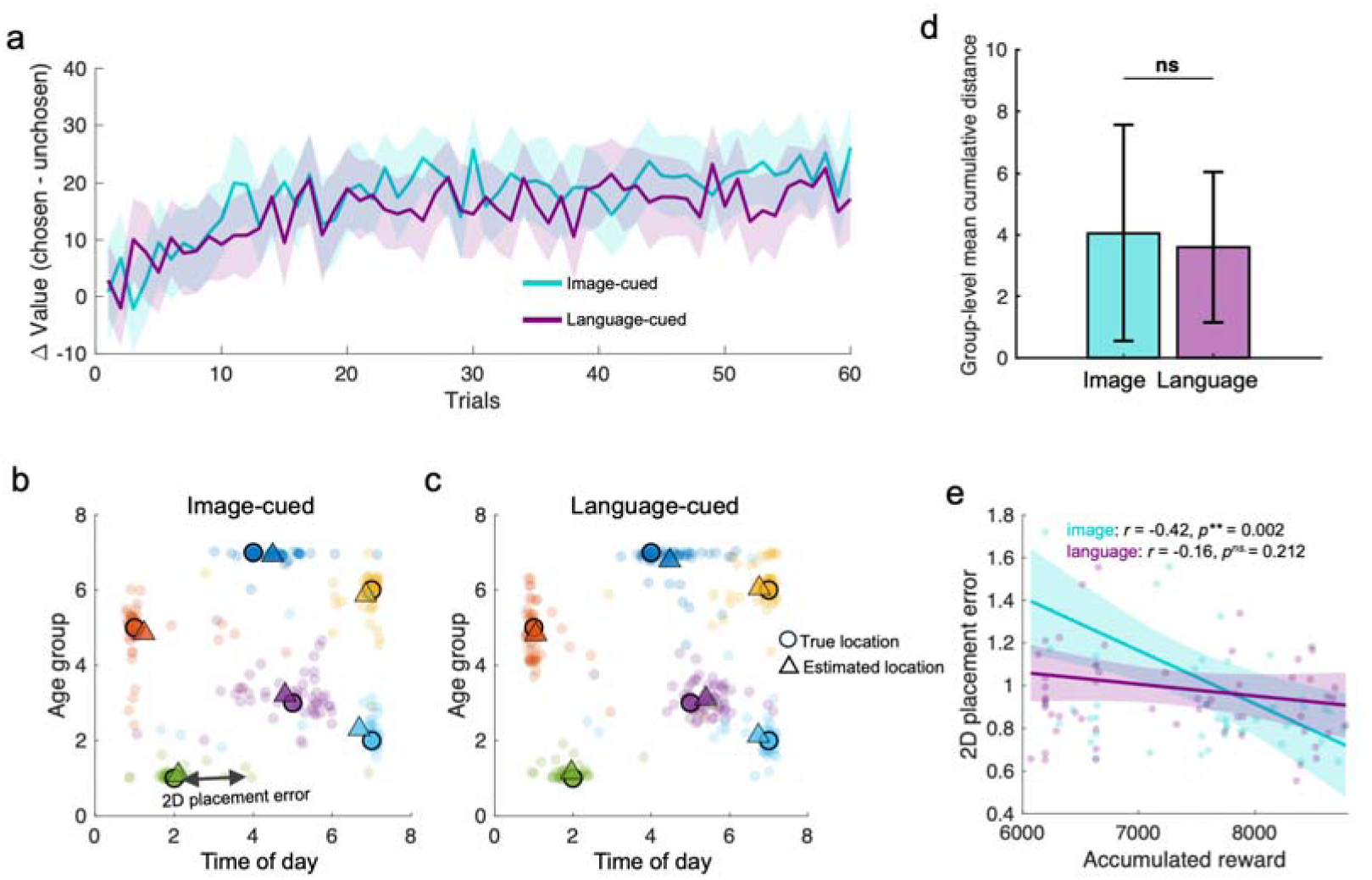
Results of the choice task. **a.** Trial-by-trial dynamics of decision value differences (Δ value = *V*_chosen_ − *V*_unchosen_) across 60 trials in two restaurant contexts for the image-cued (cyan) and language-cued (purple) conditions. Each restaurant context comprised two blocks of 30 trials; for visualization, the two blocks corresponding to the same restaurant are concatenated here. Across the experiment, the four blocks (two restaurants × two blocks) were presented in randomized order. **b-c.** Comparison of true and estimated item locations in the 2D map from the placement task for the image-cued and language-cued groups, respectively. Circles indicate the true item positions, and triangles indicate the participants’ average estimated positions. Each transparent colored dot represents one participant’s estimated location, with different colors denoting different food items (i.e., different locations). The solid arrow illustrates the 2D placement error for one participant in one example food item, i.e., the Euclidean distance between the true and estimated item locations (lower values indicate greater accuracy). **d.** *Group-level mean cumulative distance between participants’ estimated item locations and the true locations in the 2D space, shown separately for the image-cued and language-cued groups. Bars depict group means ± SEM.* **e.** Correlation between accumulated reward (x-axis) in the choice task and accumulated 2D placement errors (y-axis). The accumulated reward refers to the total gains earned during the choice task. Lines show separate linear fits for the im-age-based (cyan) and language-based (purple) conditions, with shaded areas representing 95% confidence intervals. Significance levels: \**\**p < 0.01; *ns*. = not significant.

A key question is whether more accurate spatial memory for the food items (i.e., lower 2D placement error) predicts better performance in the choice task. The depends on the condition: higher accumulated reward in the choice task was associated with lower 2D placement error in the image-cued condition (*r* = −0.42, 95%*CI* [−0.62, −0.16], *p* = 0.002), but not in language-cued condition (*r* = −0.16, 95%*CI* [−0.40, 0.09], *p* = 0.212, *BF*_01_ = 3.44; *Fis*□*er′sZ* = 1.46, *p* = 0.07; Fig. 3e), although there was no difference in accumulated reward between groups (*t* (109) = 1.00, *p* = 0.319, *BF*_01_ = 4.40).

### Higher cognitive disorganization attenuates spatial generalization specifically in visually derived cognitive maps

We compared four computational models (as specified in the pre-registration) to investigate the mechanisms underlying value-based decisions in the choice task, including a baseline model (random choice with bias), a model-free Q-learning model capturing value learning from experienced rewards, a spatial Gaussian process regression (GPR) model capturing spatial generalization, and a hybrid model combining model-free learning and spatial generalization. Full model details are provided in the Supplementary Methods.

Participants’ behavior in the choice task was best described by the Spatial-GPR model (Fig. 4a). This model assumes that reward generalization is governed by similarity between options represented in a latent 2D space, such that rewards associated with previously chosen options are weighted by their similarity to the current option, as previously applied to reward generalization in physical space^10^. In the current study, this model well predicted participants’ subjective values for the chosen food items (Fig. 4b-c; see also Supplementary Fig. S2 for posterior parameter distributions and parameter recovery).

**Figure 4:**
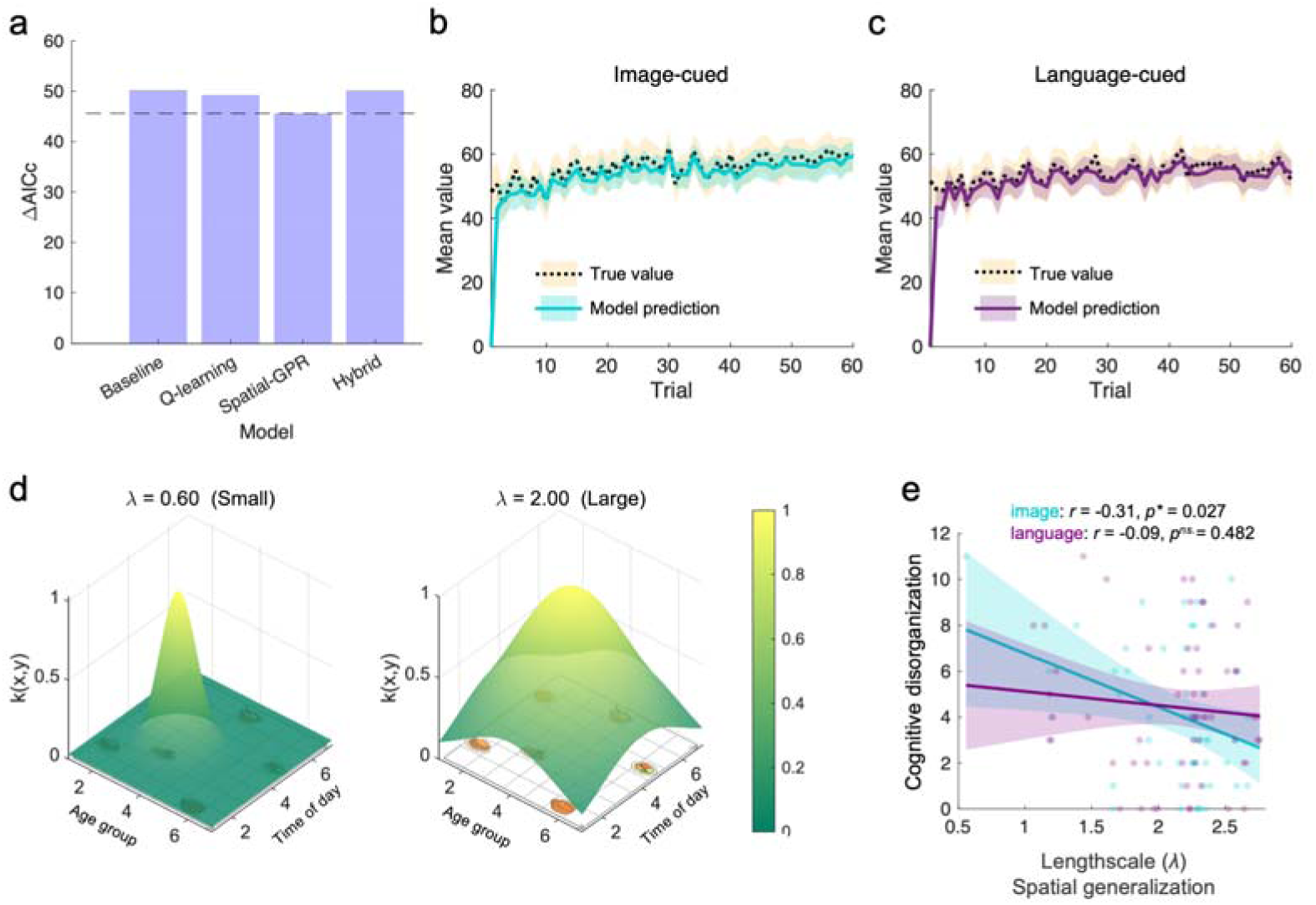
Model-based analysis of choice behavior. **a.** Model comparison based on the corrected Akaike information criterion (ΔAICc; lower values indicate better model fit). **b-c.** Comparisons of the true value (dashed line) for the chosen option with the estimated value (solid line) from the best-fitting Spatial-GPR model for the im-age-cued and language-cued conditions, respectively. The shaded areas represent 95% confidence intervals. **d.** Kernel surfaces with different length-scale parameters *λ* over a semi-transparent background image. The left panel shows a sharper surface with a small length scale (*λ*=0.60), whereas the right panel shows a broader surface with a large length scale (*λ*=2.00). Higher *λ* indicates greater spatial generalization in subjective value computations during decision-making. **e.** Correlation between the length-scale parameter *λ* from the Spatial-GPR model and cognitive disorganization score (O-LIFE subscale). Lines show separate linear fits for the image-cued (cyan) and language-cued (purple) conditions, with shaded areas representing 95% confidence intervals. Significance levels: \**p* < 0.05; *ns*. = not significant.

Under the assumption of this model, participants should be able to infer the values of items at any location on the map by generalizing from learned spatial relationships. Indeed, the Spatial-GPR model produced value predictions that were well aligned with participants’ estimated values and generalized well to food items that were never presented during the choice task (Supplementary Fig. S4). Moreover, participants whose behavior was best described by the Spatial-GPR model showed smaller deviations between model-predicted and estimated values than those described by other models (see also Supplementary Fig. S4, last panel).

This model assumes reward generalization across space via Gaussian process regression, with *λ* indexing the extent of spatial generalization, larger values indicate broader generalization (Fig. 4d). We next examined how this generalization process was related to cognitive disorganization. The results showed that narrower spatial generalization (smaller *λ*) was associated with higher cognitive disorganization. Consistent with the findings from the similarity rating task, this effect was observed only in the image-cued condition (*r* = −0.31, 95%*CI* [−0.54, −0.04], *p* = 0.027), but not in the language-cued condition (*r* = −0.09, 95%*CI* [−0.34, 0.17], *p* = 0.482, *BF*_01_ = 5.99; *Fis*□*er′sZ* = 1.18, *p* = 0.120; Fig. 4e). No associations between *λ* and other personality measures, such as the Big Five, were observed (see Supplementary Fig. S5 and Fig. S6), indicating that the observed association is selective for cognitive disorganization. Moreover, *λ* was positively correlated with mental rotation performance in the image-cued condition (*r* = 0.28, 95%*CI* [0.01, 0.52], *p* = 0.043), but not in the language-cued condition (*r* = 0.22, 95%[−0.03, 0.45], *p* = 0.084, *BF*_01_ = 1.63; see also Supplementary Fig. S6), mirroring the similarity-rating results and linking spatial generalization in the image-cued condition to individual differences in spatial transformation performance. Spatial sensitivity was weakly correlated with the GP length-scale parameter (*r* = 0.17, *p* = 0.078), suggesting that the two measures may reflect related but not identical aspects of spatial representation.

## Discussion

This study found that participants formed and used an internal 2D conceptual structure inferred from fragmented experiences, as reflected in both similarity judgments and reward-based decisions. Participants with higher cognitive disorganization showed reduced reliance on spatial representations learned from visual stimuli, whereas performance was largely preserved when the structure was provided through language. This pattern is more consistent with a difficulty in deriving coherent structure from visual perceptual input than with a general impairment in relational reasoning or reward learning. Such percept-to-structure transformations have been proposed to depend on hippocampal-entorhinal processes^23^, which may help explain why visually based mapping was more sensitive to individual differences in disorganization.

### Selective impairment for visually derived structure: links to the shallow cognitive map hypothesis

The finding that higher cognitive disorganization was accompanied by reduced reliance on spatial representations is consistent with theoretical proposals that psycho-sis-spectrum disturbances may involve instability in cognitive map representations^15^. The shallow cognitive map hypothesis proposes that psychosis involves unstable hippocampal attractor dynamics that yield weaker and less coherent relational structure. Empirical studies have reported attenuated grid-like coding during physical spatial navigation^17^, reduced global organization of semantic networks during movie watching^24^, and reduced reliance on model-based planning during reward-guided decision making^25,26^. Together, these findings suggest that schizophrenia-spectrum disorganization is associated with shallow and unstable cognitive map representations, particularly when relational structure must be inferred from visual perceptual input.

This account is also compatible with evidence that N-methyl-D-aspartate receptor hypofunction in parvalbumin positive interneurons may reduce inhibitory precision within hippocampal networks, lowering signal-to-noise ratio and destabilizing attractor dynamics that support relational binding and spatial representation^27,28^. Computational frameworks further propose that such disinhibition can yield noisier internal representations, particularly when structure must be inferred from high-dimensional perceptual input^29^, reflecting the role of latent psychological processes in shaping behavior^44^. Although the present study does not include neural measures, the selective attenuation of spatial generalization in the visually cued condition is consistent with accounts linking reduced hippocampal precision to instability in map construction.

Consistent with this interpretation, in the current study, reward-based decisions relied on the remembered spatial layout when the internal map was built from visual inputs. This pattern suggests a greater reliance on abstract spatial structure during visual-to-structure transformations; a capacity that appears to be attenuated in schizophrenia-spectrum disorganization.

Gaussian-process models have been applied to explain reward generalization in physical space^10^ and in visual feature spaces^30^. Here, we showed that reward generalization within a 2D conceptual space constructed from both image-based and language-based inputs is likewise well captured by a Gaussian-process model. We further showed that individuals with higher cognitive disorganization showed narrower generalization kernels (smaller length scales) in the image-cued condition, providing a computational signature of shallow or noisy spatial representations. Together, these findings provide a computationally grounded account that spatial sensitivity and re-ward-based spatial generalization were selectively weakened in individuals with higher schizophrenia-spectrum disorganization when structure had to be extracted from visual inputs.

### Why the association is absent in linguistic structure: evidence from language-based processing in psychosis

A striking feature of our results is the absence of schizotypy-related cognitive disorganization impairments in the language-cued condition. This dissociation provides important theoretical insight. Unlike visual stimuli, where relational structure must be actively inferred from variable sensory input, linguistic descriptions provide an explicit, symbolically structured scaffold for representing relations^31^, thereby reducing the demands on active relational integration.

Several findings from schizophrenia-spectrum research align with this interpretation. Although incoherence in language output is typically considered to reflect disorganized thinking in psychopathology^32,33^, this association is not always observed^34,35^.

Recent large-sample computational work highlights that language output anomalies can arise from multiple mechanisms, including differences in underlying associative structure, executive regulation over expression, or motivational factors, rather than providing a direct readout of thought organization^36^. Accordingly, the absence of overt linguistic incoherence does not imply intact underlying cognitive structure.

Neurocognitive studies indicate that language comprehension engages distributed semantic networks that integrate established relational knowledge accrued over the lifespan (see^37^, for a review). In contrast, processing of semantically matched pictures relies more strongly on visual-perceptual systems^38^, with word stimuli preferentially engaging linguistic and higher-level semantic regions^38^. These findings indicate that linguistic input arrives in a format that is already organized into a relational structure. In line with this view, spatial sensitivity and generalization under language input were not related to individual differences in mental spatial ability, whereas both measures covaried with mental spatial ability when inputs were image-based. This dissociation suggests that linguistic cues engage relational structures that are already organized or abstracted. These findings raise the possibility that linguistic input engages a qualitatively different mapping process from that required to infer spatial structure from perceptual experience.

Taken together, this evidence suggests that language reduces the need to derive structure from noisy perceptual input, effectively bypassing the computational step most vulnerable to cognitive disorganization. According to this view, disorganization impairs the formation of spatially coherent maps from visual cues, but not the use of structure when it is linguistically provided. This account also explains why performance in the language-cued condition did not correlate with disorganization and why cognitive tests on spatial rotation and working memory did not predict performance: linguistic descriptions do not require the construction of internal metrics in the first step.

### Conceptual and methodological contributions

This study has made several conceptual advances. First, it refines the concept of cognitive maps by demonstrating that visually derived maps and language-supported relational representations may not be equivalent, at least under the specific task demands of the present paradigm. Spatial sensitivity and generalization depend on individual differences in cognitive disorganization only when relational structure must be inferred from perceptual (image-based) experience, but not when it is accessed through linguistic cues. Second, the findings show that impairments associated with cognitive disorganization emerge selectively when cognitive map structure must be inferred and organized from visual inputs. Third, the results suggest a conceptual role for language and conceptual systems as a relatively stable relational scaffold. Linguistic input appears to provide access to relational structure that is less dependent on online spatial transformation processes, potentially buffering against vulnerabilities in map formation. This perspective offers a new angle on how language may play a protective or compensatory role within the schizophrenia spectrum.

Methodologically, the paradigm grounds abstract structure learning in meaningful, non-arbitrary associations between age, time of day, and food items, allowing relational structure to be acquired through coherent experiences. All participants remained engaged throughout the study and completed the experiment within a single session. Note that our sample was not limited to university students: 20% of participants were over 30 years of age, including a participant aged 64 years. In addition, we extend abstract knowledge maps to reward learning and decision making, and use computational modeling to show that reward generalization relies on inferred spatial structure rather than simple associative strategies.

Moreover, the current findings suggest that treatment approaches may benefit from incorporating environmental modifications that increase linguistic structure for individuals with high cognitive disorganization. In particular, explicit verbal labeling and structured psychoeducation may help make relational information more accessible, rather than requiring individuals to infer it solely from perceptual cues. However, the present study was conducted in a subclinical sample, and the cognitive disorganization measure used here should not be treated as equivalent to clinician-rated disorganization in schizophrenia.

Finally, although the present study does not include neural measures, the behavioral and computational signatures it reveals offer a powerful assay for probing distinct neural contributions to cognitive map formation and map use. This framework lays the groundwork for integrating model-based behavioral quantification with neural measures to examine how perceptual and linguistic structure interact in psychosis-spectrum cognition.

### Conclusion

Together, our findings indicate that cognitive disorganization selectively affects the construction of cognitive maps derived from visual inputs, whereas map construction under linguistic descriptions is less affected. This dissociation suggests that individual differences in schizotypal cognitive disorganization may be related to difficulties in transforming perceptual experiences into a stable relational structure. Thus, linguistic references may facilitate further cognitive processing in this condition. Understanding how perceptual and linguistic routes to structure interact, and how they fail, offers a promising path toward more mechanistic accounts of cognition in psychosis spectrum disorders. In sum, individuals with higher cognitive disorganization struggle when structure must be inferred from what they see, yet perform unaffected by cognitive disorganization when that structure is already supplied through language.

## Acknowledgements

We thank Sebastijan Veselic and Todd Hare and his research group for insightful discussions, and all participants for their contributions to this study.

## Funding

This project has received funding from the European Research Council (ERC) under the European Union’s Horizon Europe research and innovation programme (grant agreement No 101118756). It is part of the ERC Synergy project DELTA-LANG.

## Declaration of competing interest

PH has received grants and honoraria from Novartis, Lundbeck, Mepha, Janssen,

Boehringer Ingelheim, Neurolite, and OM Pharma outside of this work. No other disclosures were reported.

## Data and code availability

All data, materials, and codes are publicly available at https://github.com/xiaoyanwu2024/cogmap-ssd.git.

## Author Contributions

The study was conceptualized by X. Wu, F. Rabe, and P. Homan. Methodology was developed by X. Wu and P. Homan. Software development, formal analysis, data curation, visualization, and project administration were carried out by X. Wu. Data collection was conducted by X. Wu and P. Wu. The original draft was written by X. Wu, and the manuscript was reviewed and edited by X.Wu, Y. Pauli, S. Theves, V. Edkins, F. Rabe, W. Omlor, A. R. Misra, P. Homan, W. Hinzen and I. E. Sommer. Supervision and funding acquisition were provided by P. Homan.

## Notes

https://github.com/xiaoyanwu2024/cogmap-ssd.git

### Competing Interest Statement

The authors have declared no competing interest.

### Summary of Updates

This version has been revised in response to the reviewers' and editor's comments during the submission process.

