## Supplementary information for "Disorganization impairs cognitive maps built from visual inputs"

### Supplementary Methods

#### Behavioral analysis

To test whether similarity judgments were better explained by one-dimensional (1D) or two-dimensional (2D) representations, we compared three linear mixed-effects models that differed in how dissimilarity between stimuli was defined. In the 1D-time model, similarity was predicted by the absolute difference along the time dimension; in the 1D-age model, similarity was predicted by the absolute difference along the age dimension; and in the 2D-distance model, similarity was predicted by the Euclidean distance combining both dimensions. All models included group and its interaction with the dissimilarity predictor as fixed effects, as well as a random intercept for participant. Model fit was assessed using Akaike’s Information Criterion (AIC). Model comparison showed that the 2D-distance model provided the best fit to similarity judgments (AIC = 29,627), outperforming both the 1D-time model (AIC = 30,047) and the 1D-age model (AIC = 30,202), indicating that similarity judgments were better captured by a two-dimensional distance metric than by either dimension alone.

#### Computational modeling

##### Model description

As specified in the pre-registration, we compared four computational models with different hypotheses about decision-making during the choice task to examine how participants learned and generalized rewards across contexts. The data from all participants were included in this analysis.

*Model 1: Baseline model.* The baseline model assumes that participants make decisions with a fixed bias toward the left option, independent of previous rewards or option values. Choices were modeled with a single free parameter, 𝑝, denoting the probability of choosing the left option:

$$p(a_{t}=left)=p$$

$$p\left( a_{t}=right \right)=1-p$$

*Model 2: Q-learning model.* This model assumes that participants make decisions based on option values that are updated using a standard model-free reinforcement learning algorithm. Option values 𝑄(𝑜) are updated after each trial 𝑡 based on the received reward 𝑟𝑡 :

$$Q\left( o \right)\leftarrow Q\left( o \right)+a(rt-Q(o))$$

where 𝛼 ∈ (0, 1) is the learning rate. Choices follow a softmax rule:

$$p(a_{t}=o_{t})=\frac{1}{1+exp[\beta\left( Q\left( o_{2} \right)-Q(o_{1}) \right)]}$$

where 𝛽 is the inverse temperature controlling choice stochasticity. Two free parameters were estimated: 𝛼 and 𝛽.

*Model 3: Spatial gaussian process regression (GPR) model.* See the main text.

*Model 4: Hybrid model.* The hybrid model assumes that participants combine model-free Q-learning and model-based spatial generalization. For each option 𝑜, the hybrid value is a weighted combination of Q learning and GPR-based predictions:

$$V\left( o \right)=wQ(o)+(1-w)u(o)$$

where 𝑤 ∈ [0, 1] determines the relative reliance on Q-values versus GP predictions. Choices are generated via a softmax:

$$p(a_{t}=o_{t})=\frac{1}{1+exp[\beta\left( V\left( o_{2} \right)-V(o_{1}) \right)]}$$

The Q-values are updated after each choice using the same rule as in Model 2, and the GP memory is updated with the chosen location and reward. The free parameters are 𝛼 (learning rate), 𝛽 (inverse temperature), 𝜆 (spatial lengthscale), and 𝑤 (Q/GP weight).

##### Model fitting and comparison

Model parameters were estimated for each participant by maximizing the log-likelihood of observed choices given the model. For each participant and model, the log-likelihood is computed as

$$L=\sum_{t=1}^{T} logP(a_{t}|\theta)$$

where 𝑃(𝑎𝑡 | 𝜃) is the model-predicted probability of the observed choice on trial 𝑡, given parameters 𝜃. To reduce the risk of local minima, parameter estimation was performed in two independent runs, each using 1000 random starting points within a global optimization routine (*GlobalSearch* inMATLAB). For each subject, the best-fitting parameters were defined as those yielding the maximum log-likelihood across all runs and starting points. Model comparison was performed using the corrected Akaike information criterion (AICc), which evaluates relative model fit while penalizing model complexity and adjusting for finite sample sizes. For a model with 𝐾 free parameters, sample size 𝑛, and maximized log-likelihood log 𝐿, AICc is given by

$$AICc=-2logL\frac{2K(K+1)}{n-K-1}$$

Lower AICc values indicate better trade-offs between model fit and complexity.

##### Model validation

*Parameter recovery.* To assess parameter identifiability, we conducted a parameter recovery analysis. Synthetic choice data were generated for a subset of participants by simulating behavior from the model using their individually fitted parameters. The model was then refitted to these simulated data using the same optimization procedure as in the original analysis. For each parameter, recovered values were strongly correlated with the true generating values across participants (Supplementary Fig. S2), indicating reliable recoverability.

*Model prediction.* To evaluate the model’s predictive adequacy, we examined whether the fitted model could reproduce the trajectory of subjective value estimates observed in participants’ behavior. Using each participant’s fitted parameters, we computed trial-by-trial model-predicted expected values for the chosen option and compared these predictions with the corresponding true outcome values experienced by participants. Predicted and observed values were averaged across participants at each trial, separately for each map and cue condition, and uncertainty was quantified using 95% confidence intervals reflecting between-subject variability. Across both image-cued and language-cued conditions, the model closely tracked the evolution of experienced values over learning (Fig. 4b-c), indicating that it captured the dynamics of value updating and choice-relevant expectations.

*Value generalization.* To examine whether the model captured value generalization beyond directly experienced outcomes, we compared participants’ explicit value ratings for individual items with the model-predicted values derived from the inferred spatial map. For each participant, we used the individually estimated parameters of the spatial-GPR model to compute predicted expected values for all items in each map based on each participant’s history of experienced rewards. These model-predicted values were then aligned with participants’ rated values and compared at the item level.

We next examined value generalization at the individual level. For each participant, explicit item ratings were compared with model-predicted values derived from the fitted model, yielding participant-specific correspondence patterns. The strength of this correspondence depended on which model best accounted for the participant’s choice behavior: participants best fit by the spatial-GPR model (M3) or the hybrid model (M4) showed close alignment between rated values and model predictions, whereas this relationship was weaker for participants best fit by baseline or non-spatial learning models (M1–M2) (see Supplementary Fig. S5). Quantifying prediction error specifically for untrained items revealed that participants whose behavior was best explained by the spatial-GPR model exhibited the smallest deviations between rated and predicted values (Supplementary Fig. S5), providing direct evidence that value generalization relied on the inferred spatial structure.

### Supplementary Tables

Table S1: Demographic and psychometric measures for the image- and language-cued groups.

| Measure | Image-cued  (𝑁 = 51) | Language-cued  (𝑁 = 60) | Statistic | 𝑝 value |
| --- | --- | --- | --- | --- |
| Age (years) | 26.54 (4.66) | 27.63 (6.81) | 𝑡 = −0.96 | 0.340 |
| Gender (M/F) | 18/31 | 17/40 | 𝜒2 = 3.46 | 0.629 |
| Years of education | 16.39 (1.79) | 16.68 (2.47) | 𝑡 = −0.70 | 0.763 |
| Hunger level | 3.09 (1.60) | 3.56 (1.65) | 𝑡 = −1.49 | 0.139 |
| Minutes since last meal (min) | 234.86 (170.26) | 292.36 (279.90) | 𝑡 = −1.28 | 0.204 |
| O-LIFE total | 11.75 (6.21) | 13.33 (6.85) | 𝑡 = −1.27 | 0.207 |
| O-LIFE Unusual Experiences | 3.14 (2.48) | 3.43 (2.87) | 𝑡 = −0.58 | 0.566 |
| O-LIFE Cognitive Disorganization | 4.14 (3.09) | 4.43 (3.04) | 𝑡 = −0.51 | 0.613 |
| O-LIFE Introvertive Anhedonia | 2.67 (2.02) | 2.63 (2.09) | 𝑡 = 0.09 | 0.932 |
| O-LIFE Impulsive Nonconformity | 1.80 (1.36) | 2.83 (1.91) | 𝑡 = −3.22 | 0.002 |
| BFI Neuroticism | 23.02 (5.64) | 23.70 (6.48) | 𝑡 = −0.58 | 0.560 |
| BFI Extraversion | 25.53 (4.95) | 25.80 (5.70) | 𝑡 = −0.26 | 0.792 |
| BFI Openness | 35.94 (5.96) | 38.43 (5.53) | 𝑡 = −2.28 | 0.024 |
| BFI Agreeableness | 31.08 (3.47) | 30.55 (4.15) | 𝑡 = 0.72 | 0.473 |
| BFI Conscientiousness | 30.10 (5.67) | 29.47 (5.55) | 𝑡 = 0.59 | 0.556 |
| Positive affect | 32.10 (5.84) | 31.53 (6.42) | 𝑡 = 0.48 | 0.631 |
| Negative affect | 21.94 (6.97) | 20.58 (6.91) | 𝑡 = 1.03 | 0.306 |
| Working memory (N-back) | 0.89 (0.08) | 0.90 (0.06) | 𝑡 = −0.32 | 0.746 |
| Mental rotation task | 23.82 (7.88) | 25.72 (7.84) | 𝑡 = −1.26 | 0.209 |

Note. Values are mean (SD). BFI refers to the Big Five Inventory; O-LIFE refers to the Oxford–Liverpool Inventory of Feelings and Experiences. Positive Affect and Negative Affect were assessed using PANAS. Working memory was assessed using an N-back task, and spatial transformation ability using a mental rotation task. Bold 𝑝 values indicate 𝑝 < 0.05. All 𝑡 tests were two-tailed independent-samples tests.

Table S2: Linear mixed-effects model results predicting dissimilarity rating score.

| Fixed effects | Estimate (*β*) | SE | *t* value | *p* value | 95% CI (Lower, Upper) |
| --- | --- | --- | --- | --- | --- |
| Intercept | 1.856 | 0.385 | 4.82 | 1.442 × 10⁻⁶ | 1.102, 2.611 |
| Group (language) | 2.931 | 0.537 | 5.45 | 5.099 × 10⁻⁸ | 1.878, 3.984 |
| Distance | 0.871 | 0.048 | 18.33 | 2.984 × 10⁻⁷³ | 0.778, 0.964 |
| Cognitive disorganization (CD) | 0.401 | 0.075 | 5.36 | 8.427 × 10⁻⁸ | 0.255, 0.548 |
| Group × Distance | -0.313 | 0.066 | -4.71 | 2.542 × 10⁻⁶ | -0.443, -0.182 |
| Group × CD | -0.456 | 0.102 | -4.45 | 8.627 × 10⁻⁶ | -0.657, -0.255 |
| Distance × CD | -0.061 | 0.009 | -6.64 | 3.411 × 10⁻¹¹ | -0.079, -0.043 |
| Group × Distance × CD | 0.055 | 0.013 | 4.38 | 1.189 × 10⁻⁵ | 0.031, 0.080 |

Note. Linear mixed-effects model with random intercepts for subjects: dissimilarity ~ group × distance × CD + (1 | subid). Dissimilarity reflects reverse-scored similarity. Group was coded as image (reference) and language. Distance refers to pairwise distance in the 2D conceptual space. Cognitive disorganization (CD) was included as a continuous predictor.

### Supplementary Figures

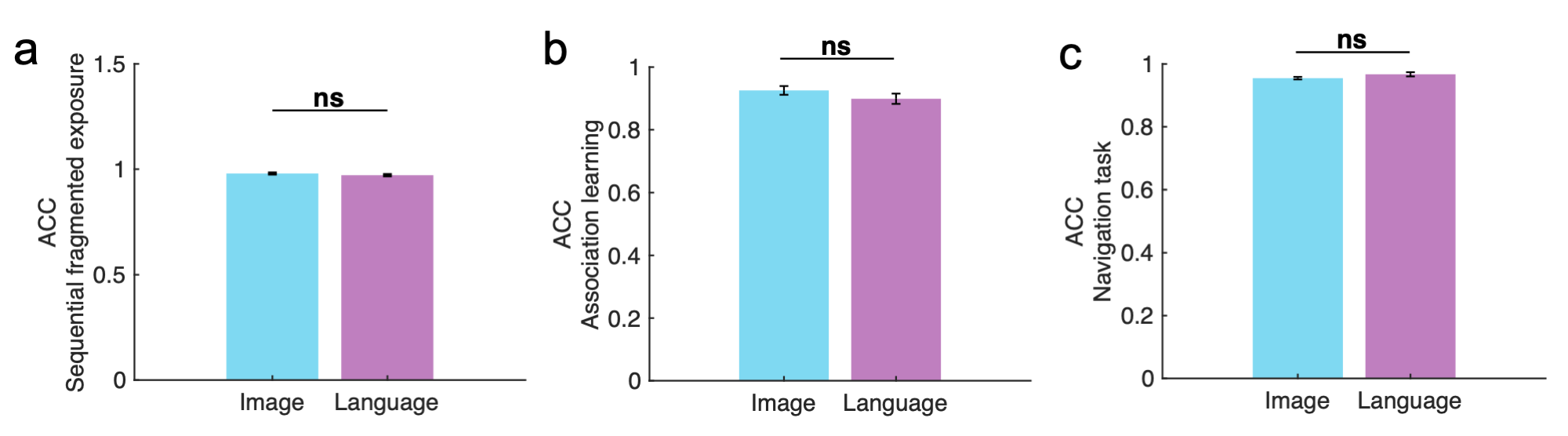

Figure S1: Group comparison of training accuracy. a. Accuracy in the sequential fragmented exposure of the ordering task. b. Accuracy in the association learning task. c. Accuracy in the navigation task. Bars show group means ± SEM. Independent-samples t-tests revealed no significant group differences (sequential fragmented exposure: 𝑡 (109) = 1.25, 𝑝 = 0.215; association learning: 𝑡 (109) = 1.21, 𝑝 = 0.228; navigation task: 𝑡 (109) = –1.50, 𝑝 = 0.137). 𝑛𝑠, not significant.

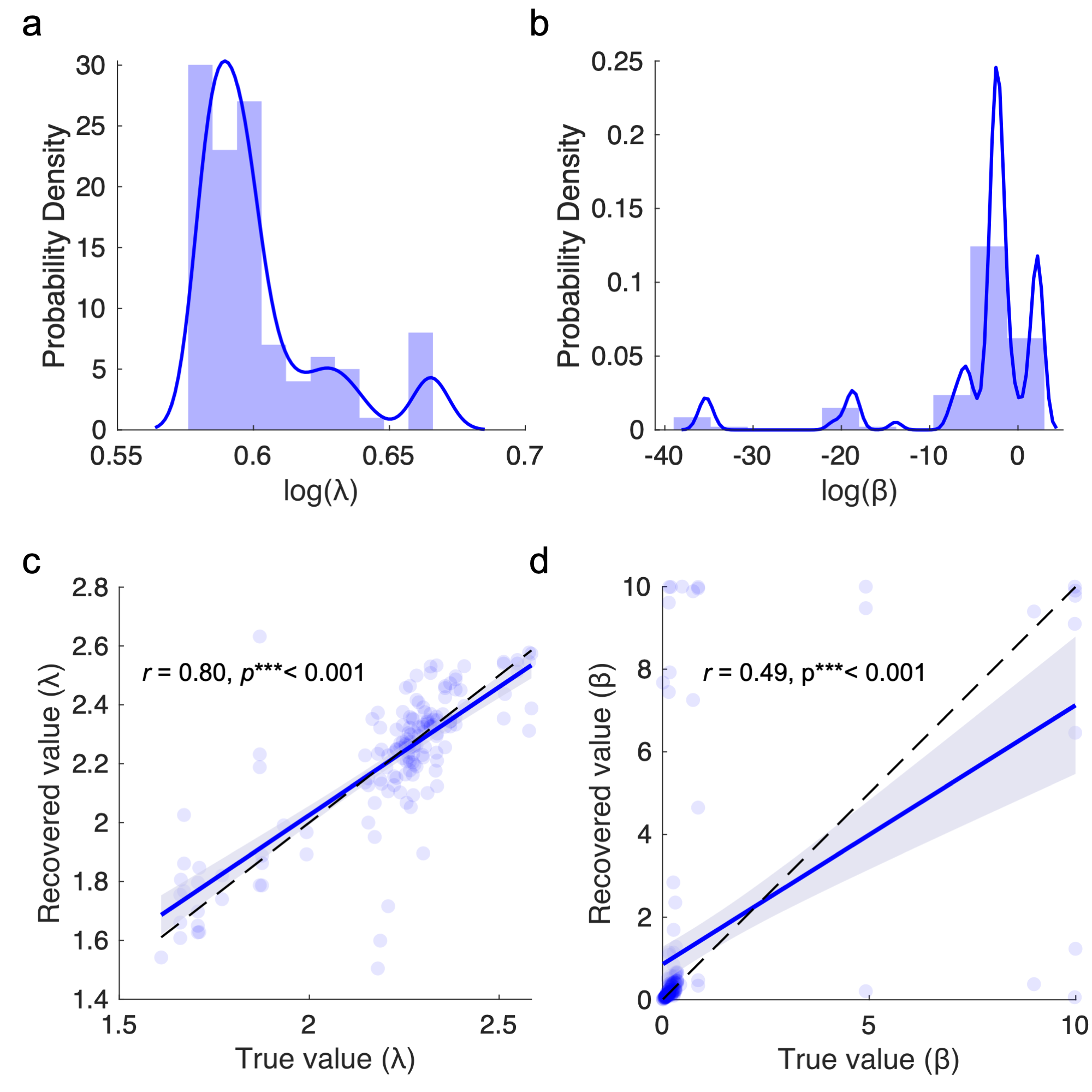

**Figure S2: Model parameter distributions and recovery. a–b.** Distributions of parameter estimates from the best-fitting model (Model 3: Spatial-GPR) across all participants. Panels display histograms with kernel density estimates for the log-transformed parameters: (a) log(𝜆) (length-scale) and (b) log(𝛽) (inverse temperature). **c–d.** Parameter recovery results for the best-fitting model (Model 3: Spatial-GPR). Panels show the relationship between true and recovered parameter values across simulated datasets for (c) the length-scale parameter (𝜆) and (d) the inverse temperature parameter (𝛽). Blue lines indicate linear fits with 95% confidence intervals, and dashed diagonal lines represent perfect recovery. Reported 𝑟 values correspond to the Pearson correlations between true and recovered parameter estimates. Significance levels: ***𝑝 < 0.001.

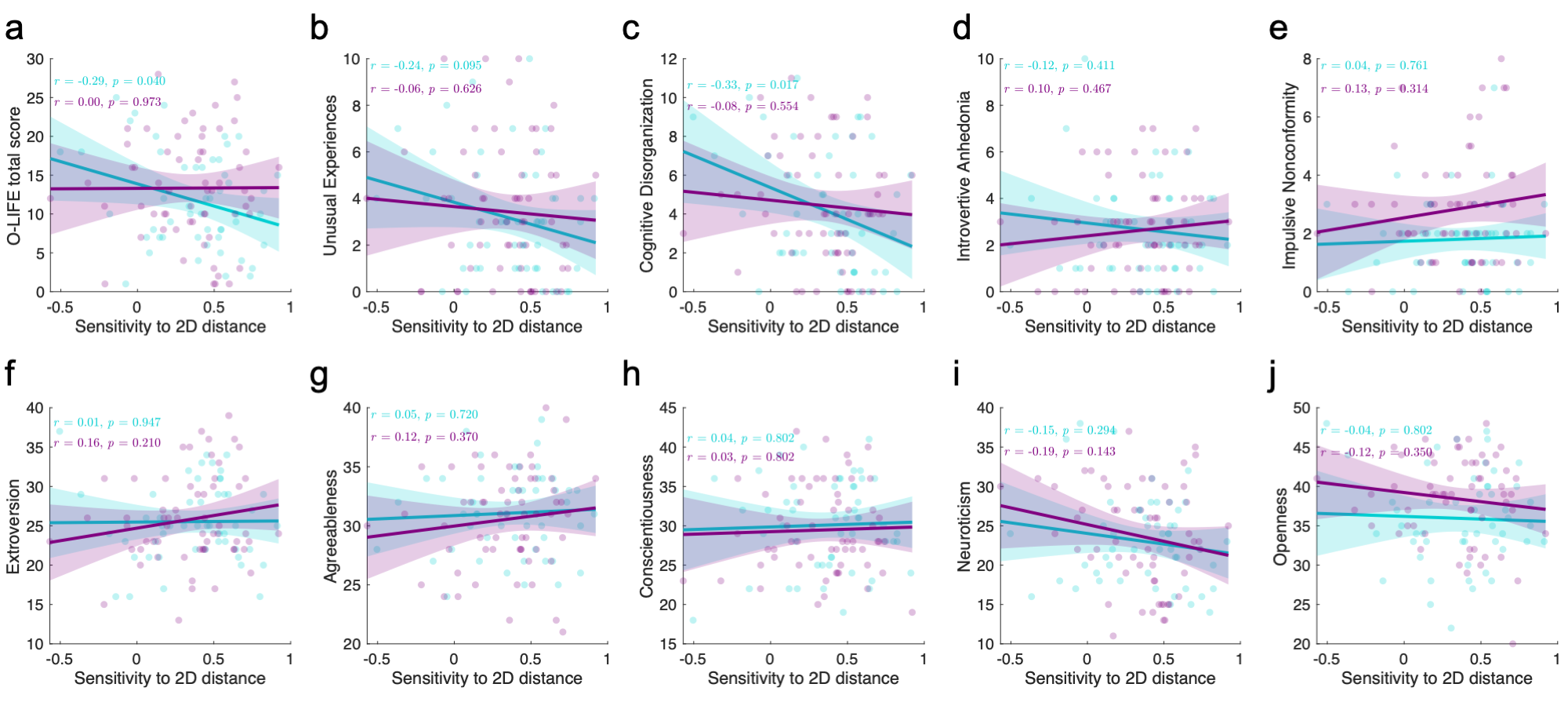

**Figure S3: Spatial sensitivity correlates with schizotypy but not Big Five traits. a–j**. Correlations between spatial sensitivity and individual difference measures. Spatial sensitivity (x-axis) reflects the Fisher 𝑍-transformed correlation between pairwise 2D distances and dissimilarity scores, with larger values indicating stronger sensitivity to spatial structure. The first row (a–e) shows correlations with schizotypy traits measured by the O-LIFE measurement, and the second row (**f–j**) shows correlations with the Big Five personality traits (control analysis). Cyan lines denote linear fits for the image-cued group and purple lines for the language-cued group, with shaded areas representing 95% confidence intervals. Reported 𝑟 values correspond to Pearson correlations.

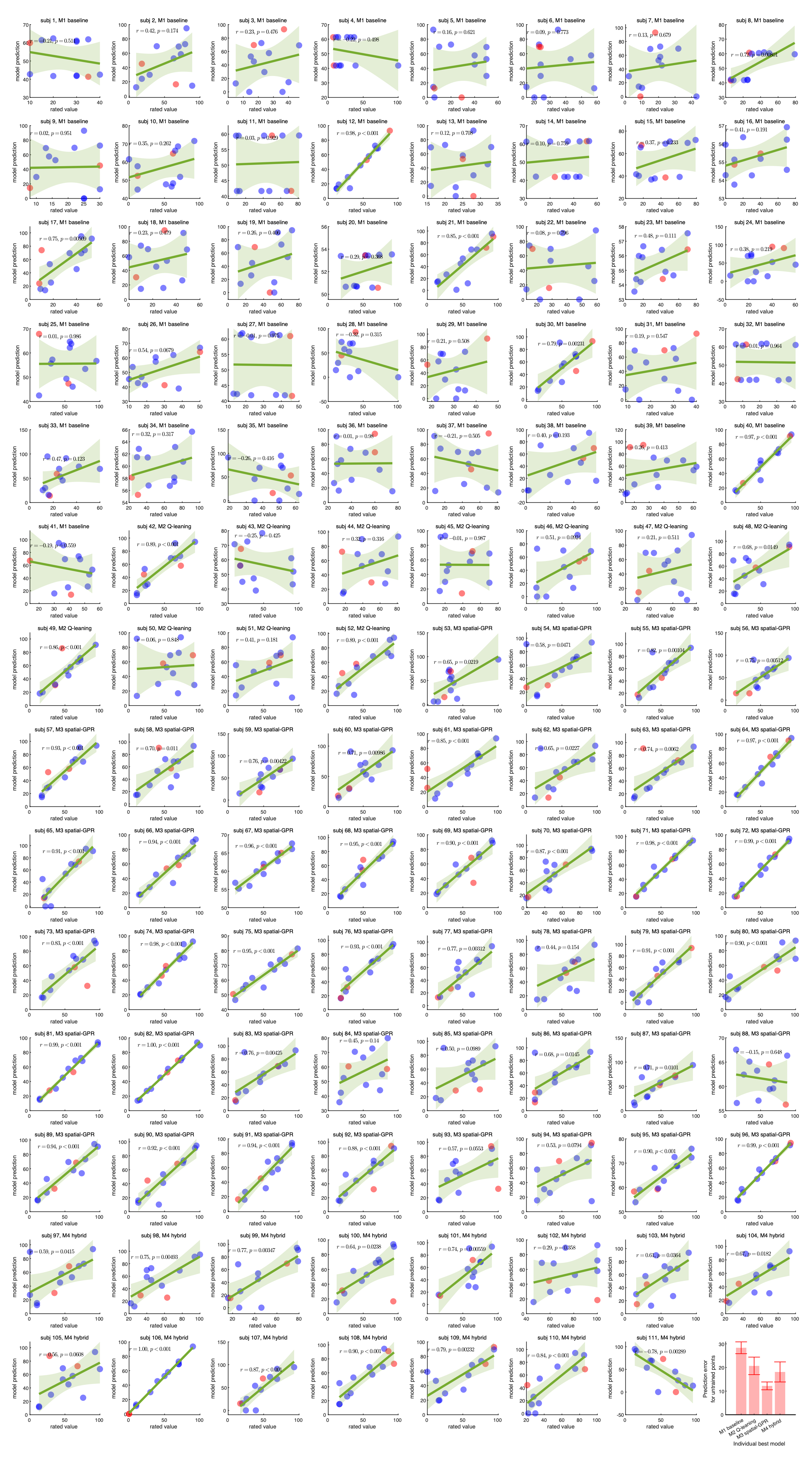

**Figure S4: Participant-level model predictions of item values.** Each panel shows the relationship between participants’ rated item values (x-axis) and model-predicted values (y-axis), computed from the group level best-fitting model (Model 3: Spatial-GPR) using participant-specific parameters as inputs. Panel titles indicate participant ID and their best-fitting model. Blue dots represent items learned during the choice task, and red dots indicate unobserved items that required value generalization (corresponding to question-mark positions in Fig.1g). Green lines denote linear fits with shaded 95% confidence intervals, and the statistics report Pearson correlation coefficients (𝑟, 𝑝). Across participants, the Spatial-GPR (Model 3) and Hybrid (Model 4) models showed stronger correspondence between predicted and rated values compared with the Baseline and Q-learning models, capturing both learned values and generalized value inference for unobserved items.

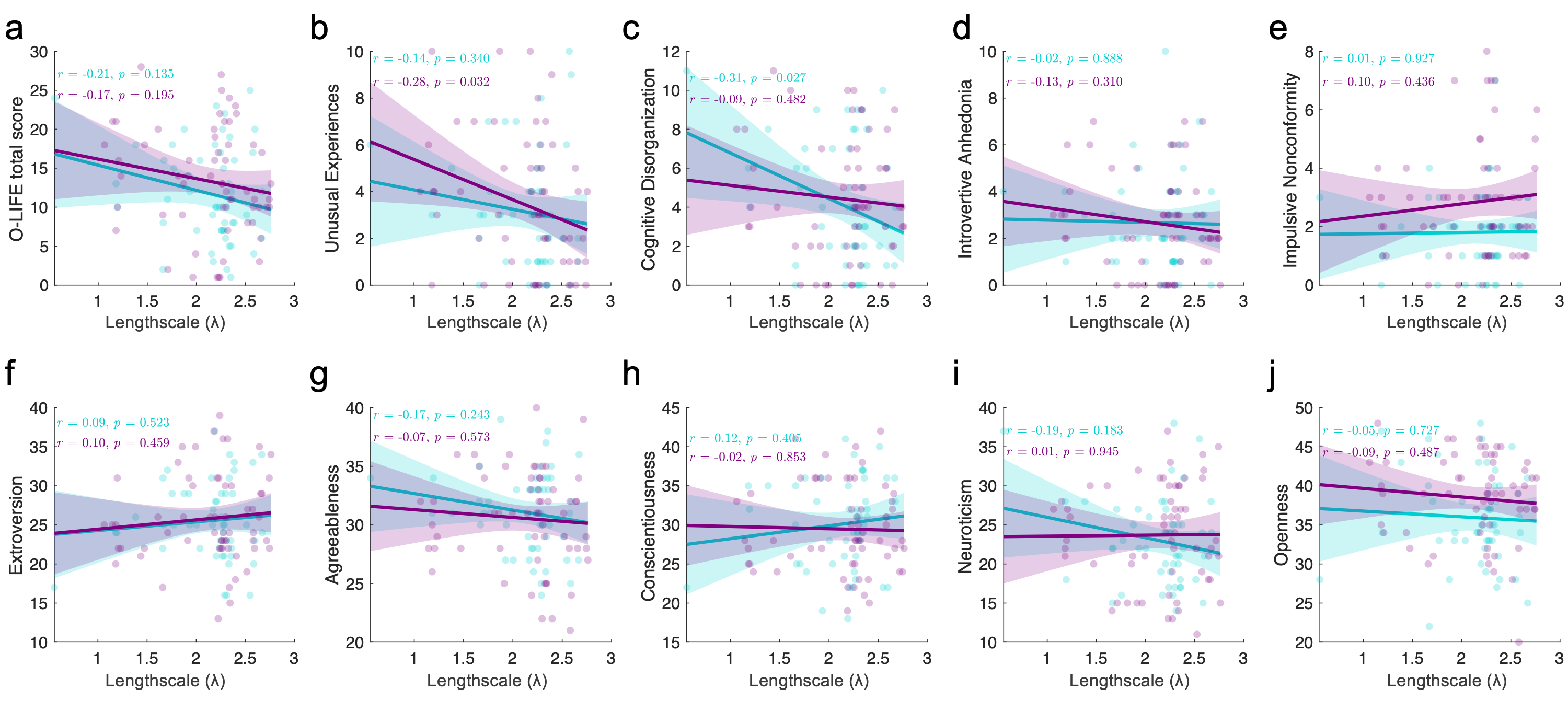

**Figure S5: Model lengthscale correlates with schizotypy but not Big Five traits.** **a–j.** Correlations between the lengthscale parameter (𝜆) from the best-fitting model and individual difference measures. The lengthscale (𝜆; x-axis) reflects the degree of spatial generalization in the model (larger values indicate broader generalization across space). The first row (a–e) shows associations with schizotypy traits measured by the O-LIFE questionnaire, and the second row (**f–j**) shows associations with the Big Five personality traits (control analysis). Cyan and purple lines indicate the image-cued and language-cued groups, respectively, with shaded areas representing 95% confidence intervals. Reported 𝑟 values denote Pearson correlations.

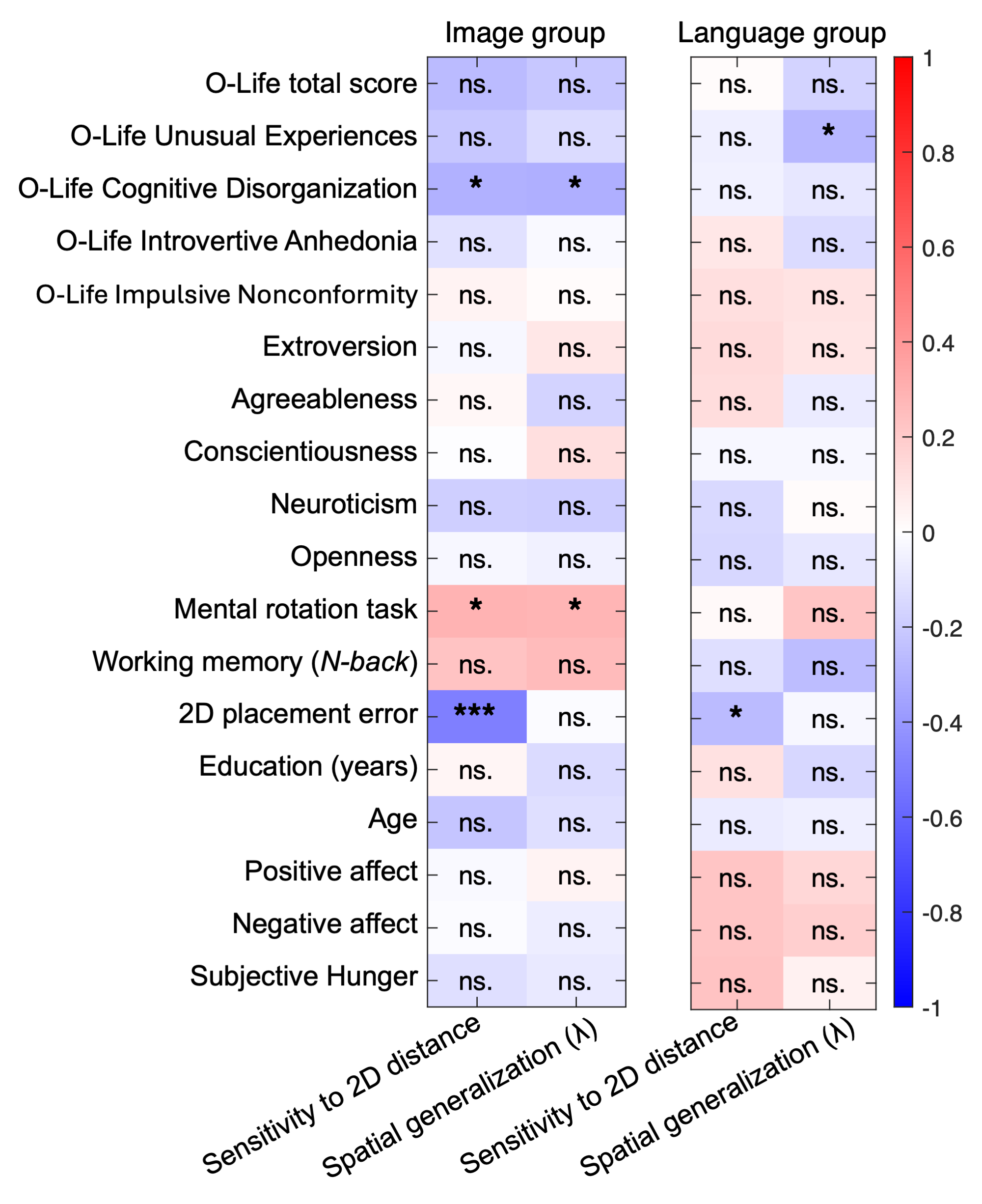

**Figure S6: Correlation matrix of psychological and behavioral measures with spatial sensitivity and generalization.** The heatmap displays Pearson correlations between individual differences (including O-LIFE subscales, Big Five personality traits, cognitive performance measures, demographic variables, and affective ratings) and two key behavioral indices: spatial sensitivity and generalization strength (𝛾). Each cell shows the correlation coefficient, color-coded from negative (blue) to positive (red) values, as indicated by the color bar on the right. Significance levels are denoted within each cell: 𝑛𝑠 = not significant; *𝑝 < 0.05; **𝑝 < 0.01; ***𝑝 < 0.001.
